# Development of a Semantically Related Emotional and Neutral Stimulus Set

**DOI:** 10.1101/2021.01.18.424707

**Authors:** Gemma E. Barnacle, Christopher R. Madan, Deborah Talmi

## Abstract

When measuring memory performance for emotional and neutral stimuli many studies are confounded by not controlling for differential semantic relatedness between stimulus sets. This could lead to the misattribution of the cause of an emotional enhancement of memory effect (EEM), because differential semantic relatedness also contributes to the EEM. Participants rated static visual emotional and neutral scenes on measures of arousal, valence, and semantic relatedness. These measures were used to create a novel stimulus set, which – in addition to demonstrating significant differences in measures of valence and arousal – also controlled for within-set semantic relatedness; thus resolving a crucial issue that has not previously been addressed in the use of visual emotional stimuli. As an added advantage, the stimulus set developed here are controlled for measures of objective visual complexity, also implicated as confounding to the investigation of memory. This article introduces a collection of emotional and neutral colour images which can be organised flexibly according to experimental requirements. These stimuli are made freely available for non-commercial use within the scientific community.

## 1. Introduction

Dimensional models of emotion suggest that emotions can be classified using two dimensions: arousal and valence (Bradley, Greenwald, Petry, & Lang, 1992; Rubin & Talarico, 2009; Russell, 1980; Watson & Tellegen, 1985). Accordingly, emotional stimulus sets typically consist of emotional stimuli which vary on arousal and are either positive or negative in valence; whereas neutral stimuli typically demonstrate a low level of arousal, and are rated as neither positive nor negative (e.g., International Affective Picture System [IAPS]: Lang, Bradley, & Cuthbert, 1997, 2008). These two dimensions of arousal and valence reliably predict aspects of the emotional experience associated with viewing visual stimuli, such as autonomic responses, and have been used extensively to reliably manipulate emotion in experimental settings.

While many valenced pictorial stimulus sets are already available (e.g., International Affective Picture System [IAPS]: Lang, Bradley, & Cuthbert, 1997, 2008; Nencki Affective Picture System [NAPS]: Marchewka, Zurawski, Jednoróg, & Grabowska, 2013; Geneva Affective Picture Database [GAPED]: Dan-Glauser & Scherer, 2011), none control for the semantic relatedness or objective visual complexity of the stimuli, which are important because they may influence the dependent measure. By controlling these factors and providing the raw values for these measures, we aim to advance the field into line with the quality typical for word stimuli sets, where ratings for many stimulus dimensions have been investigated (e.g., Larsen, Mercer, & Balota, 2006). For example, the MRC Psycholinguistic Database provides ratings for up to 26 factors for several thousand words (Coltheart, 2007), and software such as the Latent Semantic Analysis (LSA) can provide a measure of the semantic cohesion of word lists (Landauer, Foltz, & Laham, 1998; Landauer & Dumais, 1997; Dillon, Cooper, Grent-’t-Jong, Woldorv, & LaBar, 2006; Shiffrin, 2003).

### 1.1 Semantic relatedness as a confound to empirical psychological investigation

Based on Tulving’s (1979) definition of semantics as understanding or knowledge, we use the term *semantic relatedness* to refer to a shared meaning among stimuli; which has been operationalised in a number of different ways (Buchanan, Westbury, & Burgess, 2001; Griffiths, Steyvers, & Tenenbaum, 2007; Hunt & McDaniel, 1993; Nelson, Kitto, Galea, McEvoy, & Bruza, 2013). The measurement of semantic relatedness has also been investigated in a variety of ways. For example, early investigations used category membership (Cohen, Bousfield & Whitmarsh, 1957; Battig & Montague, 1968; Battig & Montague, 1969, Van Overschelde, Rawson, & Dunlosky, 2004), where participants were given a category title e.g. “seafood” and were asked to write down as many items that should be included in that category as possible. Pair matching procedures have also been employed, in which participants are presented with all possible pairs of stimuli and asked to rate the relatedness of the pairs (e.g. for words: Talmi & Moscovitch, 2004; and pictures: Talmi, Schimmack, Paterson, & Moscovitch, 2007a). Previous studies have also investigated the semantic relatedness of words by using computational methods to estimate cases of word-context congruence and incongruence, which together give a reliable estimate of the semantic relatedness of two or more words (Landauer et al., 1998).

For the current work, it was important to draw attention to *within-set* semantic relatedness, which we define as a measure of how well each stimulus of a set represents the given theme, averaged across the stimuli of that set. The within-set semantic relatedness of negative emotional stimuli is often high – many stimuli convey a shared meaning or share a common theme such as war, poverty, and violence. Similarly, these related pictures are more likely to share other properties such as the complexity of the scene (Ochsner, 2000), the presence of people (Talmi, Luk, McGarry, & Moscovitch, 2007) and certain hues such as red (due to many pictures showing bloody scenes). Conversely, neutral stimuli are generally more heterogeneous as they often do not share a common theme. When investigated empirically, it has been demonstrated that this differential level of semantic relatedness between emotional and neutral stimuli embodies a confound which unduly influences the measure of subsequent memory. One example comes from the emotional enhancement of memory (EEM) literature (Talmi, 2013; Talmi & Moscovitch, 2004). Talmi and Moscovitch (2004) showed participants semantically related emotional, semantically related neutral, and semantically *unrelated* neutral words in discrete lists and measured participant’s free recall memory after ~45 minutes. Memory performance for semantically related emotional words was greater than that of semantically unrelated neutral words (resulting in an EEM effect); however, when semantically related emotional words were compared to semantically related neutral words the participant’s memory performance was equivalent (i.e. the EEM effect disappeared when lists were matched for semantic relatedness).

In light of this evidence and others (e.g. Balconi & Ferrari, 2012; Buchanan, Etzel, Adolphs, & Tranel, 2006; Madan, Caplan, Lau, & Fujiwara, 2012; Schwarze, Bingel, & Sommer, 2012; Siddiqui & Unsworth, 2011; Sommer, Gläscher, Moritz, & Büchel, 2008; Talmi, Luk, McGarry, & Moscovitch, 2007; Talmi, Schimmack, Paterson, & Moscovitch, 2007b) showing the contribution of semantic relatedness to EEM for both verbal and pictorial stimuli; and in light of many studies referring to this effect by way of interpretation of their findings without explicit use of semantically related stimulus sets (Kalpouzos, Fischer, Rieckmann, MacDonald, & Bäckman, 2012; Madan, Fujiwara, Caplan, & Sommer, 2017; Maratos, Allan, & Rugg, 2000; Verde, Stone, Hatch, & Schnall, 2010; Wang & Fu, 2011) the need for a stimulus set controlling for this confound is evident. This is the primary aim of the present study.

### 1.2 Visual complexity as a confound to empirical psychological investigation

In addition to semantic relatedness, other dimensions that may vary between stimulus sets should also be considered. One such dimension is visual complexity. Indeed, several studies using emotional images have attempted to control for variability in visual complexity (e.g., Kensinger, Garoff-Eaton, & Schacter, 2007; Kensinger, Piguet, Krendl, & Corkin, 2005; Ochsner, 2000; Sakaki, Niki, & Mather, 2012; Talmi, Luk, et al., 2007b). This can be particularly important since visual complexity has been shown to influence memory (e.g. Isola, Xiao, Parikh, Torralba, & Oliva, 2013; Nguyen & McDaniel, 2014). As findings suggest that subjective ratings of visual complexity (Madan, Bayer, Gamer, Lonsdorf, & Sommer, 2018) and vividness (Todd, Talmi, Schmitz, Susskind, & Anderson, 2012) can be biased by emotion, it is preferable to use computational methods to objectively measure visual complexity. Here we operationalize visual complexity using three objective, computational metrics: edge density (amount of image that is detected as ‘edge’), feature congestion (clustering of visual details), and subband entropy (disorganisation in the image, inversely related to spatial repetitions) (Madan et al., 2018; Rosenholtz, Li, & Nakano, 2007).

## 2. Method

### 2.1 Participants

Thirty-three undergraduate Psychology students from the University of Manchester were invited to take part in this study for course credit. One participant was excluded from all analyses due to not following task instructions. Four additional participants did not complete Task 2 (relatedness task) due to taking longer breaks and not finishing the tasks in time. Thus, analyses of Task 2 comprised of 28 participants (mean age = 19 years, 2 identified as males, the other as females), and analyses of Task 1 (emotion task) comprised of 32 participants (mean age = 19 years, 4 identified as males). The number of participants was determined on the basis of a previous study (Talmi, Luk, McGarry, & Moscovitch, 2007), where N=12 was sufficient to achieve reliable ratings. Note that picture scores and the comparison between picture sets compare stimulus scores that are averaged across participants.

This study was approved by the University Research Ethics Committee at the University of Manchester. Due to the high number of emotional stimuli presented in this experiment a distress policy was developed for use in this study. The function of this policy was to monitor the progression of participants through the tasks of the experiment; to monitor the mood of participants, and to act as an indicator for the reporting adverse events (policy available upon request). As part of this policy, participants made subjective ratings on four measures using 9-point scales of: Bored-Engaged, Unhappy-Happy, Anxious-Calm, Miserable-Cheerful – before and after the experiment. Adverse events were defined as a) situations where testing was curtailed due to participant distress, or b) where the participant completed testing but indicated that their mood was lower than their first rating *and* at floor (levels 1 – 2 on the scale). In addition to the above, an outcome was defined as an adverse event only when their mood did not improve after following the actions listed in the distress policy. No adverse events were recorded during this experiment.

### 2.2. Materials

786 colour pictures (size 280 x 210 pixels) of equal numbers emotional (*n*=384) and neutral (*n*=384) scenes were selected using Google Images™ and supplemented with images from the IAPS (Lang, Bradley, & Cuthbert, 1997, 2008). Pictures were cropped to size 280 x 210 pixels. Many of the selected stimuli were images with no copyright restrictions. In accordance with UK copyright law, the stimuli which were copyrighted may be used here and re-used when intended for non-commercial purposes. Due to the volume of stimuli, we are not able to acknowledge the sources of all stimuli.

The emotional scenes were negatively valenced, arousing emotional scenes. These emotional scenes were chosen exclusively because the dominant modulation model of EEM posits that arousal drives the effect (McGaugh, 2004), rather than valence being the most important factor. Recent human models of EEM (Mather, Clewett, Sakaki, & Harley, 2016) also propose a central role for arousal rather than valence. In addition to this, evidence from neuroimaging studies suggests that there may be a brain basis for selecting only negative stimuli: that negative stimuli elicit ‘stronger’ (greater amplitude) and/or more reliable activations in the brain. For example, electroencephalography (EEG) studies suggest that negatively valenced, emotional stimuli elicit a greater amplitude electrophysiological response than do positively valenced emotional stimuli (Dolcos & Cabeza, 2002; Kaestner & Polich, 2011); especially, but not always, when investigating the differential emotional electrophysiological correlates of subsequent memory (Olofsson, Nordin, Sequeira, & Polich, 2008; Righi et al., 2012; Schupp, Flaisch, Stockburger, & Junghöfer, 2006). Moreover, a recent meta-analysis (Lindquist, Satpute, Wager, Weber, & Barrett, 2015) investigating neuroimaging evidence for the representation of emotion in the brain found that although a network of brain areas could be identified as responding to emotion per se (i.e. both negative and positive in valence), it was found that negative stimuli more frequently elicited activity in some of those areas compared to positive stimuli. However, none of those areas were more frquently activated by positive compared to negative stimuli. Finally, many studies investigating emotion have often used only negatively valenced stimuli, rather than both negative and positive emotional stimuli (e.g. Maratos et al., 2000; Pottage & Schaefer, 2012; Schaefer et al., 2002; Schwarze et al., 2012; Talmi, Luk, et al., 2007; Talmi & McGarry, 2012; Taylor et al., 1998; Todd, Evans, Morris, Lewis, & Taylor, 2011; Watts, Buratto, Brotherhood, Barnacle, & Schaefer, 2014), suggesting that a set containing only negatively valenced emotional stimuli would serve the needs of researchers in this field.

Neutral scenes were selected based on the theme ‘domestic scenes’ following the previous work of our lab (e.g. Talmi, Schimmack, et al., 2007). Choosing a theme such as this allowed the selection of a large number of stimuli whilst maintaining the probability of semantic relatedness. Neutral images were selected to be complex scenes and most depicted people. For example, most stimuli contained a person carrying out an action; and plain pictures of objects (e.g., a hammer on a white background) were intentionally *not* selected. This general primary selection criteria made it more likely that neutral and emotional stimuli could later be matched on their average ratings such as visual complexity.

Within each set of emotional and neutral pictures 50% of the images formed “group A” and 50% formed “group B”. Stimuli from group A and B were ‘content-matched’ but not identical. For example if one neutral group A picture depicted a person laying paving, the corresponding neutral group B picture would also contain a non-identical picture showing a similar scene of a person laying paving. Content-matched pictures could be given the same title but were not identical (as in Kensinger et al., 2007). Pictures were selected in this manner to allow future experiments to use content-matched pictures. For example, this may be useful to match targets and lures in recognition memory tests or in ‘same-similar’ paradigms (Kensinger, Garoff-Eaton, & Schacter, 2006; Kensinger et al., 2007).

Nine neutral sub-categories which embodied the theme domestic scenes were defined as follows: Indoor “Do It Yourself” (D.I.Y); Outdoor D.I.Y; cleaning and chores; leisure scenes; hobbies and games (non-sport); hobbies and games (sporting); gardening; personal hygiene; and working at home. For the emotional scenes, nine sub-categories were defined as follows: law enforcement and armed services; children in danger; injured and wounded; medical scenes; torture; aggravated crime; poverty; death; and accidents, emergencies, and disasters. One picture representing each of these emotional and neutral sub-categories was randomly selected to be used as the ‘example matrix’ in Task 2, therefore leaving 768 experimental pictures.

### 2.3. Procedure

Each participant took part in two tasks, which we called the emotion task and the relatedness task. In the emotion task participants viewed 768 randomly ordered emotional and neutral pictures displayed on a computer screen one at a time. Participants were asked to rate each picture for emotional valence (first) and arousal (second) using computerised SAM scales (Bradley & Lang, 1994) presented on-screen, one above the other. The SAM scales have 9 intervals each and range from −4 = ‘unhappy’ to +4 = ‘happy’ for the valence scale, and −4 = ‘excited’ to +4 = ‘calm’ for the arousal scale (a central interval, ‘0’ marks the mid-point on both scales). This task was self-paced and participants were encouraged to take breaks if needed.

The relatedness task comprised of two parts, reflecting the rating of the emotional and neutral stimuli separately (using the same procedure), with the order counterbalanced across participants. When rating a given set of stimuli (emotional or neutral) participants saw the corresponding example matrix as detailed above, and this remained on the screen throughout the task to serve as an anchor. Participants were first instructed to study the aforementioned nine example pictures of the current set. Participants were advised that the examples shared a common theme or meaning, and were asked to look at the example pictures and think about what this shared meaning may be. After scrutinising the nine examples, participants were instructed to rate 422 randomly ordered experimental pictures one at a time using a 7-point Likert scale of relatedness, with 1 titled ‘low relatedness’ and 7 titled ‘high relatedness’ (as in Talmi & McGarry, 2012; see Figure 1) to judge if the trial image was semantically related to the nine examples. The 422 pictures comprised 384 of the same valence as the example matrix (emotional or neutral), and 38 were from the other valence set. The latter were included as ‘catch trials’ to ensure that participants were paying attention throughout the experiment. This task was also self-paced and participants were encouraged to take breaks if needed. After participants completed the relatedness ratings of one valence category (e.g. emotional stimuli), the procedure was repeated with the other set (e.g. neutral stimuli). On average participants took a total of two hours to complete all tasks. This experiment was realised using Cogent 2000 (Wellcome Department of Imaging Neuroscience, UCL, UK; http://www.vislab.ucl.ac.uk/cogent_2000.php). The instructions to participants were as follows:

> Screen 1:
>
> *“In this task you will help us to validate a stimuli set by providing ratings for possible pictures to be included. Nine EXAMPLE pictures of one stimulus set will be displayed throughout the experiment. TRIAL pictures will be presented one at a time and you will be asked to rate how related is the TRIAL to the EXAMPLES. The next screen explains the ratings. Press SPACE BAR to continue.”*

> Screen 2:
>
> *“When you rate the relatedness of the TRIAL to the EXAMPLES we want you to focus on whether the overall meaning of the TRIAL and the EXAMPLES are related, i.e. do they convey a similar meaning or a different meaning? We DO NOT want you to consider low-level similarities, such as colours, shapes, number of objects / people. You will rate the relatedness using a scale of 1-7 which will be displayed on the screen for each trial. Press SPACE BAR to continue.”*

**Figure 1.**
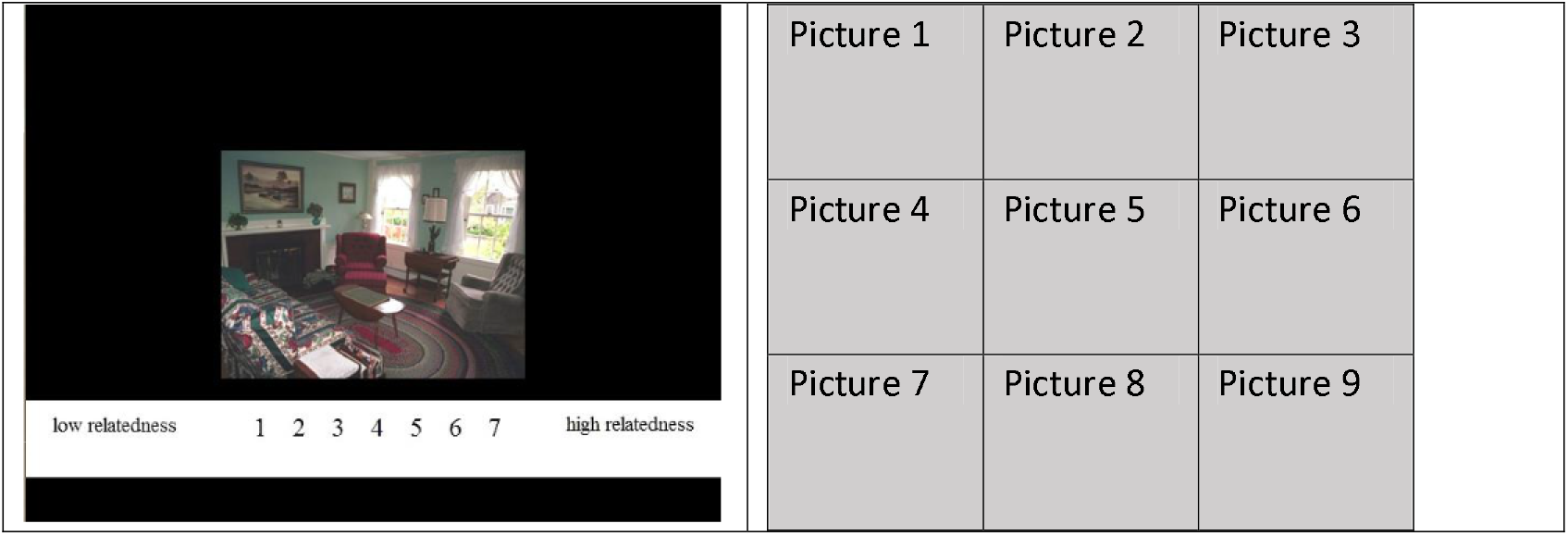
Example trial rating of semantic relatedness as seen by participants. Left: Trial image and relatedness scale. The scale remained on the screen throughout the trial. Right: Nine example images (neutral example matrix). The examples remained on screen throughout the task.

The final screen gave an example of the relatedness scale (see Figure 1).

### 2.4. Analysis

For ease of reference we converted the scores of the SAM scales to a scale of 1 – 9; where for the valence scale the converted score of ‘1’ corresponded to the SAM scale score of ‘-4’ (unhappy) and the converted score of ‘9’ corresponded to the score of to ‘+4’ (happy). For the arousal scale a converted score of ‘1’ corresponded to ‘+4’ (calm) and a converted score of ‘9’ corresponded to a SAM scale score of ‘-4’ (excited). For both converted scales, a score of ‘5’ represented the mid-point.

Responses from each participant for the emotion task (*n*=32), and the relatedness task (*n*=28) were averaged across participants for each picture. To determine the reliability of the measures used, Cronbach alpha scores were calculated. Reliability was high for all subjective measures: arousal α=.99; valence α=.98; relatedness α=.99. For analysis purposes we used mean scores of arousal, valence, and semantic relatedness, averaged across participants for each stimulus (in line with previous reporting of statistical testing of novel stimuli sets e.g. Dan-Glauser & Scherer, 2011; Lang et al., 1997; Marchewka et al., 2013).

Emotional pictures were removed from analysis if their average arousal score was less than 6, their average valence score was less than 4, and their average relatedness was less than 4 (for the relatedness ratings scale the score of ‘4’ represented the mid-point). Neutral items were removed if their average arousal score was greater than 4, their average valence score was less than 4 or greater than 6, and their average relatedness score was less than 4.

Visual complexity was calculated using three computational methods taken from Rosenholtz et al. (2007): edge density, feature congestion, and subband entropy. Edge density is based on identifying boundaries between objects and features within an image. Here we converted the images to CIELab 1976 colour space (designed to mimic the responses of the human eye) and then computed the Canny edge detection on the L* dimension using the lower and upper thresholds. (0.11 and 0.27, respectively). Feature congestion quantifies how ‘cluttered’ an image is and incorporates colour, luminance contrast, and orientation. Subband entropy quantifies the ‘disorganisation’ within the image, through Shannon’s entropy in spatial repetitions of hue, luminance, and size (i.e., spatial frequency)..

Stimuli were removed from further analyses if they were identified as outliers on any one (or more) of the measures: arousal, valence, semantic relatedness, edge density, feature congestion, or subband entropy. Outliers were identified by any score greater than or less than 2.5 standard deviations from the mean, following Thompson (2006). A quality control procedure was also carried out, whereby any two remaining stimuli from the same set that were deemed too alike but that were not intentional duplicates were identified and one stimulus removed. We also removed stimuli which were deemed to be of a poor quality (e.g., due to obvious pixelation or distortions). In order to obtain the desired outcomes – namely statistically significant differences between emotional and neutral items for arousal and valence, and no statistical differences between emotional and neutral items for semantic relatedness and visual complexity measures – we removed the 25 neutral stimuli with highest subband entropy measures; and 25 further neutral stimuli whose semantic relatedness measures were highest.

The remaining stimuli were then organised into two final sets, Emotional and Neutral (see Figure 2 for flow diagram of final stimulus selection). Given the close matching of these stimuli within sets, we anticipate that researchers would choose subsets of these final stimuli according to experimental need. The mean scores of all ratings per stimulus are provided in the supplementary materials.

**Figure 2.**
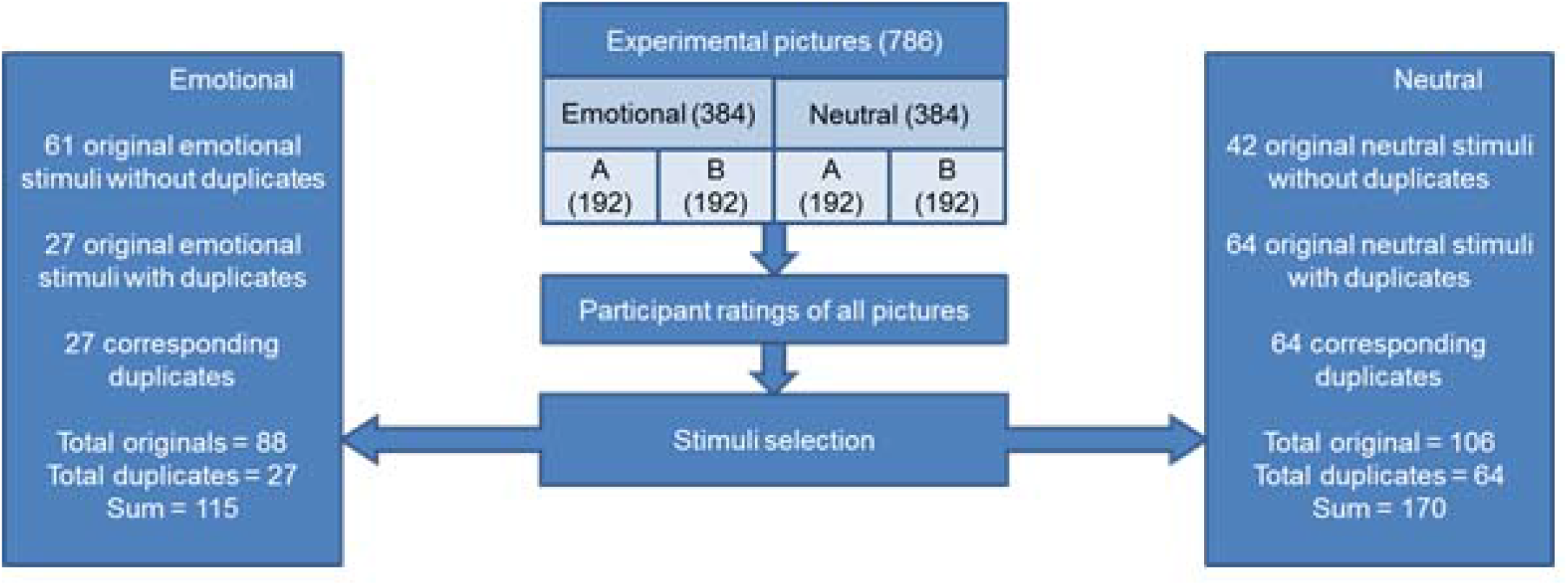
Diagram showing process of stimulus set creation. See Analysis section for detailed description of the stimulus selection. Stimuli from group A and B were ‘content-matched’ but not identical.

After stimulus selection the number of remaining emotional stimuli from either group A or B was 88, and these stimuli formed part of the final Emotional stimuli set. For clarity, these pictures will be referred to as ‘Originals’ to distinguish them from their corresponding content-matched stimuli referred to as ‘Duplicates’. Of these Original stimuli, 27 were determined to have a corresponding content-matched stimulus which also passed the selection procedure. Therefore, the final Emotional set contains 115 emotional stimuli (88 Originals and 27 Duplicates). Of the Emotional Originals, 22 were taken from the IAPS (Lang, Bradley, & Cuthbert, 1997, 2008, IAPS 2205, 2800, 3015, 3030, 3051, 3102, 3110, 3120, 3170, 3266, 3500, 3550, 6212, 6315, 6550, 9042, 9250, 9253, 9400, 9420, 9433, 9921). ^1^ The related_pictures_database (other than the IAPS) and the related_pictures_ratings are available as supplementary data.

The number of remaining neutral stimuli from either group A or B was 106, and these stimuli therefore formed part of the final Neutral stimulus set. Of these stimuli, 64 had a corresponding content-matched stimulus which passed the selection procedure. Therefore the final Neutral set contains 170 neutral stimuli (106 Originals and 64 Duplicates). Table 1 provides grand average arousal, valence, and semantic relatedness scores (averages were generated across participants per stimuli, and then averaged across stimuli to provide one average score for each measure per valence, and per set); plus objective measures of visual complexity (as described above).

**Table 1.**
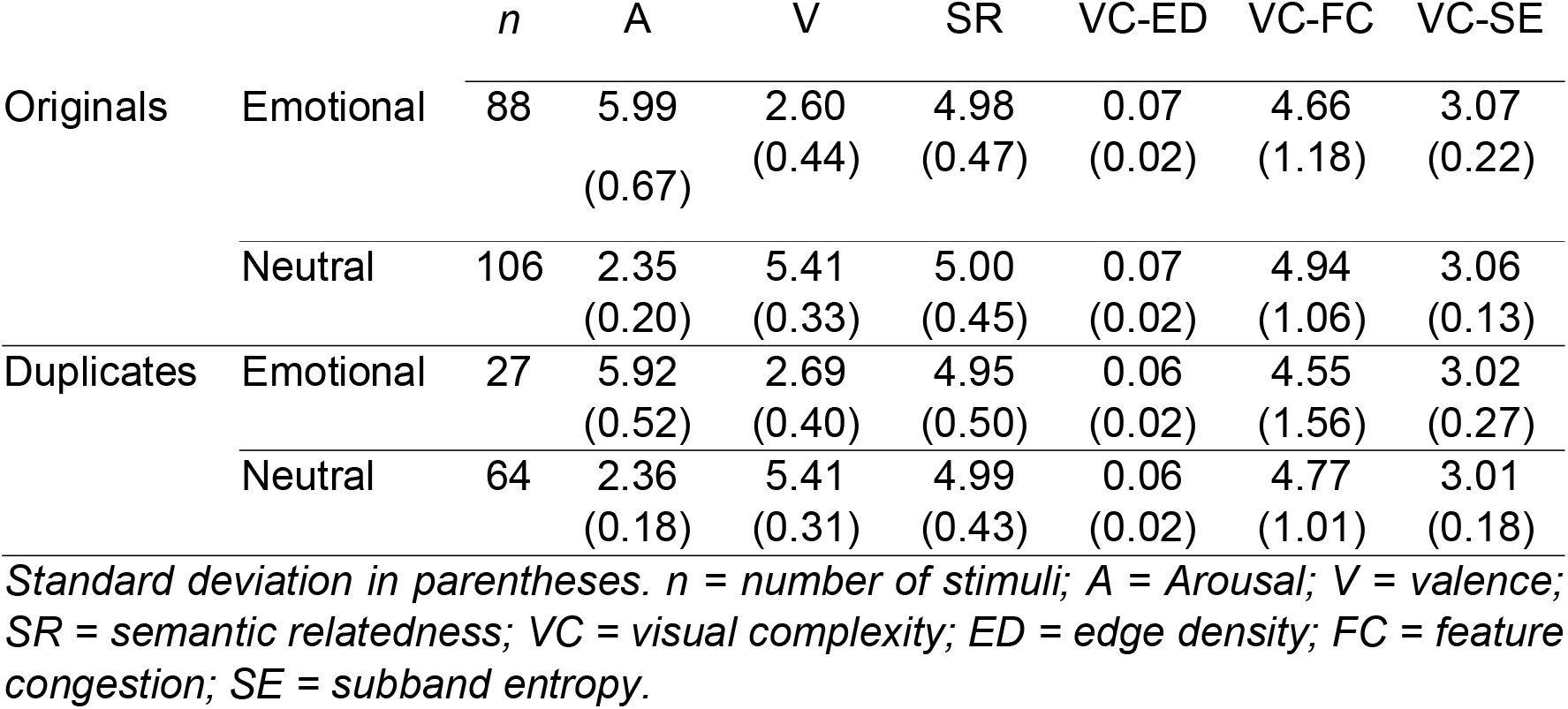
Mean and standard deviation values for arousal, valence, semantic relatedness, and visual complexity measures per valence and set for the final images.

To determine whether there existed a significant difference in the measures of arousal, valence, semantic relatedness, and objective measures of visual complexity between emotional and neutral stimuli, we conducted independent-samples t-tests, which treated the stimuli as cases. Analyses were performed separately for Originals and Duplicates. Consistent with reporting of new stimuli sets in the literature (Dan-Glauser & Scherer, 2011; Lang et al., 1997; Marchewka et al., 2013) we also represent the pictures in arousal-valence space (see Figure 3), and test the correlation of arousal and valence using the Pearson correlation coefficient.

**Figure 3.**
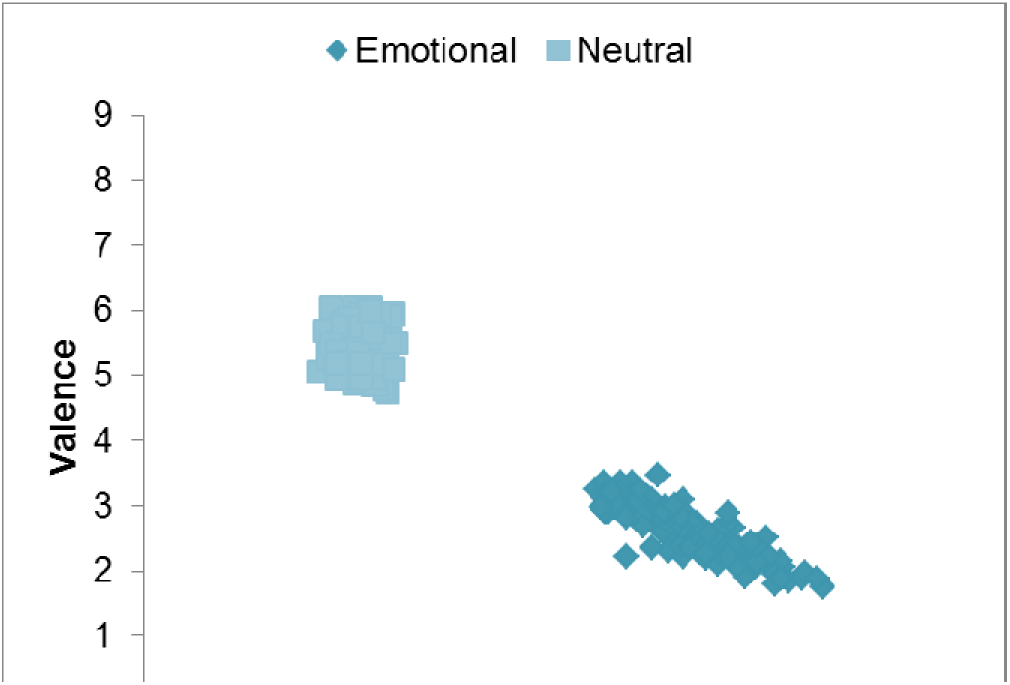
Plotting the SeRENS Emotional and SeRENS Neutral in Arousal-Valence Space.

## 3. Results

### 3.1 Originals

Analysis comparing the original emotional (*n*=88) and neutral (*n*=106) stimuli revealed a significant difference in the average arousal score. Emotional stimuli measured higher on arousal (*M* = 5.98, *SD* = 0.67) than neutral stimuli (*M* = 2.35, *SD* = 0.20), *t*(192) = 48.97, *p* < .001, 95% CI [3.48, 3.77], Cohen’s ds = 7.63. Analysis also identified a significant difference in the average valence scores of emotional and neutral stimuli. Emotional stimuli measured lower on valence, i.e. were more negatively valenced (*M* = 2.59, *SD* = 0.44) than neutral stimuli (*M* = 5.41, *SD* = 0.33), *t*(192) = 49.15, *p* < .001, 95% CI [−2.93, −2.71], Cohen’s ds = 7.35 (effect sizes calculated according to resources from Lakens, 2013). As expected, there was no significant difference in the average relatedness scores of emotional and neutral stimuli. There was also no significant difference between emotional and neutral stimuli for the visual complexity measures.

### 3.2. Duplicates

Analysis comparing the Duplicate emotional (*n* = 27) and Duplicate neutral stimuli (*n* = 64) again revealed a significant difference in the average arousal scores. Emotional stimuli measured higher on arousal (*M* = 5.92, *SD* = 0.53) than neutral stimuli (*M* = 2.35, *SD* = 0.18), *t*(89) = 34.32, *p* < .001, 95% CI [3.36, 3.77], Cohen’s ds = 10.98. Emotional stimuli measured lower on valence, i.e. were more negatively valenced (*M* = 2.69, *SD* = 0.41) than neutral stimuli (*M* = 5.41, *SD* = 0.32), *t*(89) = 31.25, *p* < .001, 95% CI [−2.90, −2.56], Cohen’s d_s_ = 7.96. As intended, there was again no significant difference in the average relatedness scores of emotional and neutral stimuli, and no significant differences for any of the measures of visual complexity.

To ensure that the corresponding original and duplicate stimuli were adequately matched we performed independent-samples t-tests for all measures comparing only the Original and corresponding Duplicates within each valence set (i.e. comparing the 27 Emotional Originals with their corresponding 27 Emotional Duplicates; and 64 Neutral Originals with their corresponding 64 Neutral Duplicates. Consistent with adequately matched stimuli, we found no significant differences between Originals and Duplicates for emotional or for neutral stimuli on any measures.

We also checked that participants could identify the ‘catch trials’ (different-valence trials) in the relatedness task; and as expected we found that these trials were reported as significantly less related to the example matrices than the same-valence stimuli (emotional stimuli: *t*(420) = 12.39, *p* < .001, Hedges’s g_s_ = 2.09; neutral stimuli: *t*(420) = 26.76, *p* < .001, Hedges’s g_s_ = 4.54).

### 3.3 Correlations

In line with publications introducing other affective stimuli sets (Lang et al., 1997; Marchewka et al., 2013) we also investigated the correlation of arousal and valence for all stimuli (averaged over participants) within Emotional and Neutral (including Duplicates). For the Emotional pictures we found a significant negative correlation between these two measures *r* = −.862, *p* = <.001, demonstrating that as ratings of arousal increased, so too did ratings of valence decrease (i.e arousing stimuli were rated as more negative in valence). For the Neutral pictures there was no significant correlation between the measures of arousal and valence *r* = .094, *p* = .22 (see Figure 3).

## 4. Discussion

The final emotional stimulus selection comprise stimuli of high arousal, negatively valenced, colour pictures; with the final neutral stimulus selection comprising low arousal, neutral valence, colour pictures; and all sets have high within-set semantic relatedness. By controlling for semantic relatedness and three objective measures of visual complexity, we can assert that any differences in behavioural response (and/or other physiological measures) to Emotional and Neutral pictures are not confounded by these measures; allowing researchers more confidence that any differences are due to emotion – the manipulation of interest. The present set of pictures also convey an advantage over existing stimuli sets as they are supplied with a number of duplicates, which may prove useful for example when testing memory using recognition paradigms.

### 4.1 Correlation of Arousal and Valence

As demonstrated in Figure 3 and evidenced by correlational analysis, the Emotional pictures exhibited a significant negative correlation between the measures of valence and arousal. That is to say, that like other stimulus sets (Marchewka et al., 2013), as the average rating of valence for a given stimulus decreased (i.e., the stimulus was perceived as more negative), the average rating of arousal for the same picture increased (i.e., the stimulus was perceived as more arousing). Like the NAPS (Marchewka et al., 2013), the present pictures demonstrated a linear relationship between valence and arousal which is different to the ‘boomerang-shaped’ pattern which represents the correlation of arousal and valence for the IAPS (Lang et al., 1997). This difference is due to the fact that our picture set contains no positively valenced stimuli (or in the case of Marchewka et al., 2013; no positive and arousing stimuli). Another difference between the spread of arousal and valence for this compared to other published stimuli sets such as IAPS (Lang et al., 1997), the NAPS (Marchewka et al., 2013), and the GAPED (Dan-Glauser & Scherer, 2011) is that the parcellation in arousal and valence space for each set is more distinct here, namely, the limits of arousal and valence are not overlapping between emotional and neutral stimuli. This may be advantageous in some studies because it suggests that based on the arousal and valence scores, the stimuli are well-categorised as either emotional or neutral. This means that we can be reasonably confident that no emotional stimulus would be incorrectly identified as neutral and vice versa, which should translate to more reliable results of emotional manipulation in future experiments. For other studies, however, this may present a disadvantage.

### 4.2 Methodological Evaluation

The objective measures of visual complexity used here were deemed to be optimal in comparison to subjective ratings of visual complexity, which is susceptible to cognitive and emotional biases. This was demonstrated by Madan et al. (2018) which showed that subjective ratings of visual complexity correlated with arousal and valence ratings, while measures of objective visual complexity did not. Though other objective measures of visual complexity have also been used (e.g., JPEG file size), the measures used here were preferred as they are thought to better model processes shown to occur in early visual cortices and have been shown to capture more inter-item variability (Madan et al., 2018). As visual complexity has been found to influence memory (e.g., Isola et al., 2013; Nguyen & McDaniel, 2015), but is often considered to be orthogonal to emotion, it is preferable to minimize this additional source of variability, as we have here. By providing stimuli with additional experimental controls, we hope to improve the precision and robustness of future studies.

Given the unsuitability of previous methods such as computational methods popular for assessing the semantic relatedness of words, and which require common contexts (LSA, Landauer & Foltz, 1998); and pair matching (Talmi, Schimmack, et al., 2007a) – which would have resulted in an inordinate number of trials (over a billion ratings for the current sets) we created an alternative novel method to measure semantic relatedness. This method was intentionally designed such that in the final sets no single stimulus or specific combination of stimuli was crucial for the integrity of the within-set semantic relatedness of each set. A weakness of our method was that because every stimulus was judged for its relatedness to a matrix of exemplars for that set, we cannot ascertain that all stimuli judged to be highly related to the exemplars would necessarily be highly related to each other. For example, two stimuli may have been rated as highly related to the exemplars because they were highly related to non-overlapping aspects of the exemplar pictures. Therefore, we expect that users of the present database of pictures would wish to verify that their subsets are comparable in relatedness using the same method described here. Such future work is advised to include a balance between genders, because as is common in psychology courses, our participants included few participants who identified as male.

The method we used employed a unipolar Likert scale of semantic relatedness ranging from 1-7 (as in Talmi & McGarry, 2012) allowed participants to be specific in their responses, whilst avoiding an overwhelming large or obviously contrived scale (as every same-set picture was designed to be semantically related to the example set on which their rating was based). Being unipolar, the scale created a ‘forced choice’ situation – i.e. there was no option to respond ‘not sure’ or don’t know’ (which participants often mistake as the midpoint of bi-polar scales), ensuring all stimuli were accurately rated.

### 4.3 Flexibility of Stimulus Selection for Future Studies

In contrast to the stringent matching process undertaken to create a well-controlled database of picture, there remains an important flexibility of stimulus selection for future studies, which represents a considerable advantage. This flexibility is due to the provision of ratings data (see supplementary materials), and the provision of duplicates. The supplementary materials provide the average arousal, valence, semantic relatedness, and three visual complexity measures for each individual stimulus. This allows researchers to hand-pick a subset of stimuli from our set according to experimental need, and to verify the statistical significance of important measures.

A further benefit of our picture set is the provision of content-matched duplicates. As well as ensuring no significant differences between originals and corresponding duplicates, we have verified that the emotional and neutral duplicates differ statistically on measures of arousal and valence, but not on semantic relatedness or measures of visual complexity – following the same pattern of significant differences as the original sets. Duplicates will be useful in future research which aims to test memory using recognition paradigms, for example where participants would be presented with a mixture of “old” (seen before) stimuli which they are required to distinguish from “new” (previously unseen) stimuli. Using content-matched lures presents an advantage over randomly selected lures because they test a higher level of accuracy of recognition, ensuring that the participant recognises a stimulus specifically, over and above the vague content of (or a familiarity with-) the stimulus. Of course, the pictures could be used without the duplicates, for example, in experiments which test memory using other methods such as free recall.

Finally, it is noted that the experience of emotion is a complex and multifaceted phenomena which challenges scientific investigation. The definition of emotion has been contested (Izard, 2010), and a number of theoretical frameworks compete to describe it (Damasio, Everitt, & Bishop, 2007; Ekman, 1992; Lazarus, 1991; Russell, 1980). Recent compelling evidence advocating the existence of mixed emotions (feeling two opposite emotions at the same time; Berrios, Totterdell, & Kellett, 2015) also pose a challenge to these models. As such there are many potential influences of emotional experience and emotional memory, and therefore these stimuli sets are limited in that they address a finite number of contributory factors. The pictures provided here are limited in that they only include highly arousing, negative pictures, and excluding either low-arousing negative pictures or positive pictures. Providing these novel, modern, validated picture stimulus sets will expand the choice of valenced pictorial stimuli; and should encourage experimenters to control for differential semantic relatedness and visual complexity in future research.

### 4.4. Conclusion

Our experiment resulted in two stimulus sets that will be optimal for use in experiments where otherwise differential semantic relatedness and/or visual complexity may prove confounding to the interpretation of the independent measure, such as the measurement of memory performance (Talmi & Moscovitch, 2004; Isola et al., 2013). As far as we are aware this is the only stimuli set of this nature, and therefore is likely to serve as an important resource for researchers of emotional memory.

## Supporting information

related_pictures_ratings

related_picture_database

## Acknowledgements and Funding Information

This study was supported by funding from the Economic and Social Research Council [ES/J500094/1]. The authors would like to thank J Hanratty for her contribution to the study design; and A Valji and AC Brebenar for their contributions to stimuli selection.

## Footnotes

1 These pictures are not supplied as part of the database. Permission should be sought from http://csea.phhp.ufl.edu/media/iapsmessage.html in order to use these stimuli, if required.

