## Supplementary figures and images for "Development of a Semantically Related Emotional and Neutral Stimulus Set"

### neutral01a.bmp

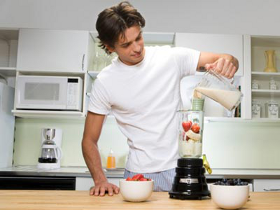

### neutral01b.bmp

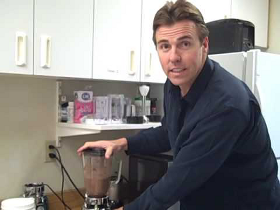

### neutral02a.bmp

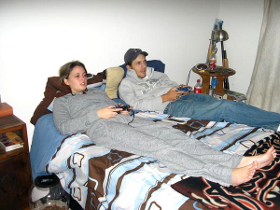

### neutral02b.bmp

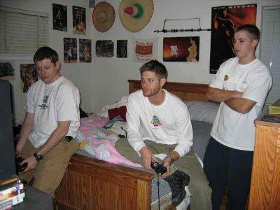

### neutral03a.bmp

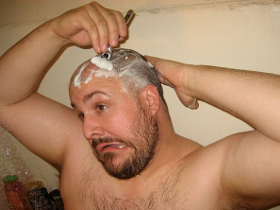

### neutral03b.bmp

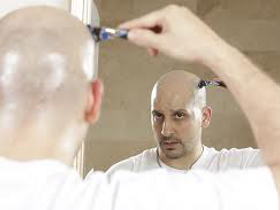

### neutral04a.bmp

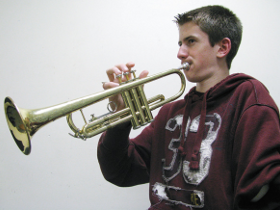

### neutral04b.bmp

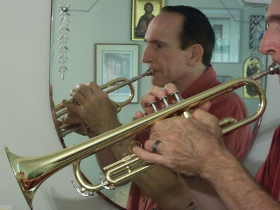

### neutral010b.bmp

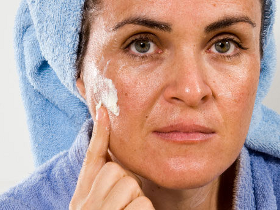

### neutral011a.bmp

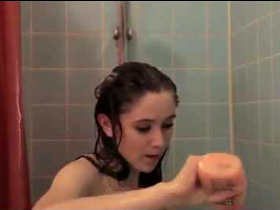

### neutral011b.bmp

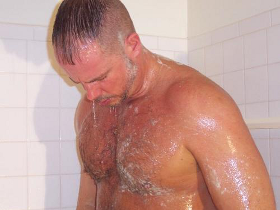

### neutral012a.bmp

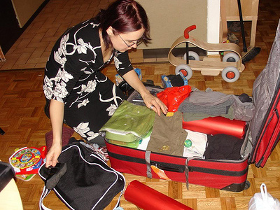

### neutral012b.bmp

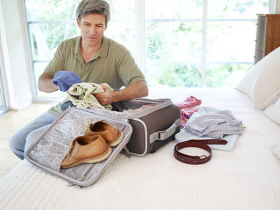

### neutral013a.bmp

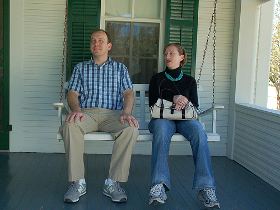

### neutral013b.bmp

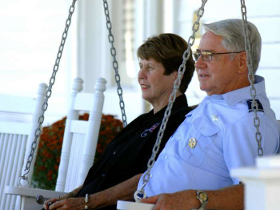

### neutral014a.bmp

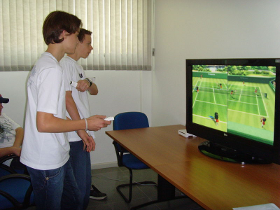

### neutral014b.bmp

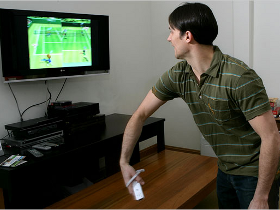

### neutral015a.bmp

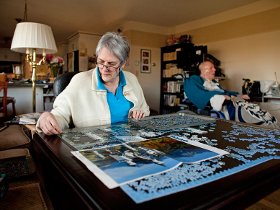

### neutral015b.bmp

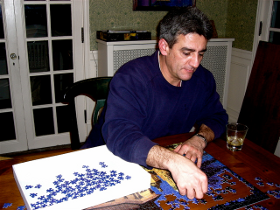

### neutral016a.bmp

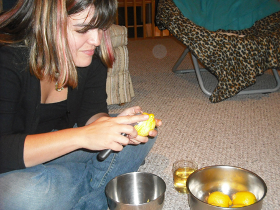

### neutral016b.bmp

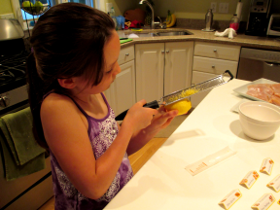

### neutral017a.bmp

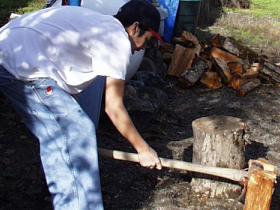

### neutral017b.bmp

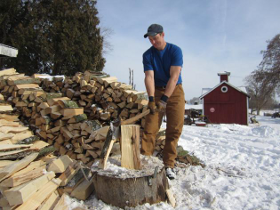

### neutral018a.bmp

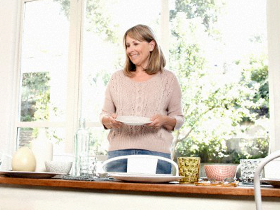

### neutral018b.bmp

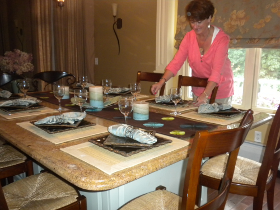

### neutral019a.bmp

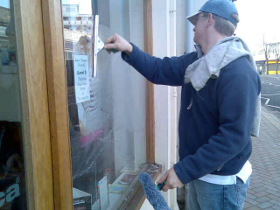

### neutral019b.bmp

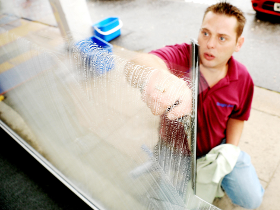

### neutral020a.bmp

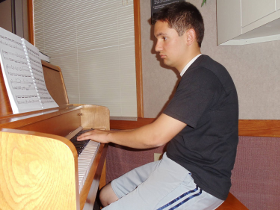

### neutral020b.bmp

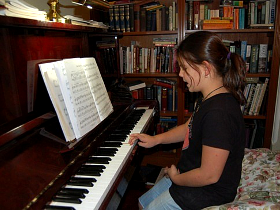

### neutral021a.bmp

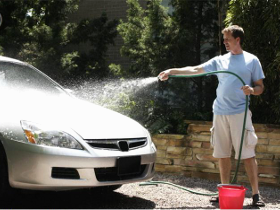

### neutral021b.bmp

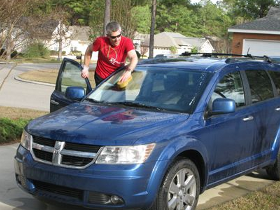

### neutral022a.bmp

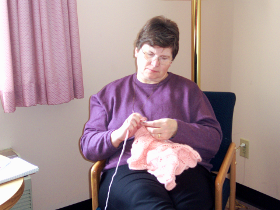

### neutral022b.bmp

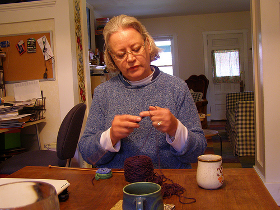

### neutral023a.bmp

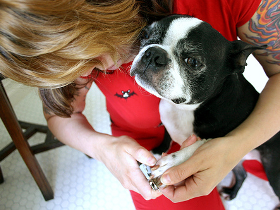

### neutral023b.bmp

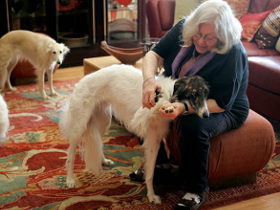

### neutral024a.bmp

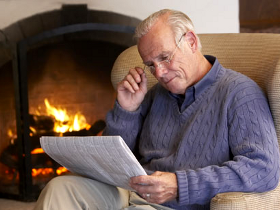

### neutral024b.bmp

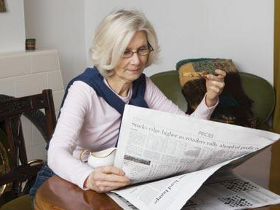

### neutral025a.bmp

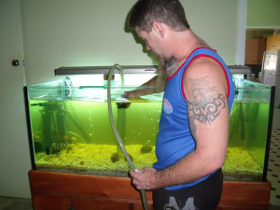

### neutral025b.bmp

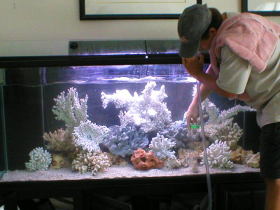

### neutral026a.bmp

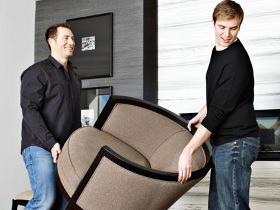

### neutral026b.bmp

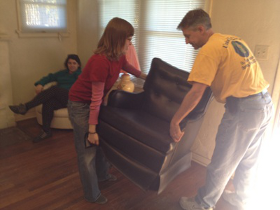

### neutral027a.bmp

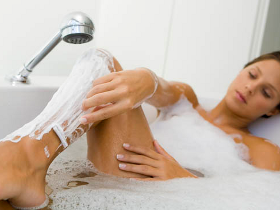

### neutral027b.bmp

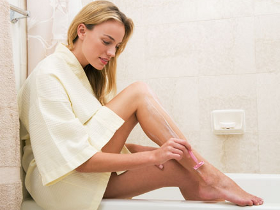

### neutral028a.bmp

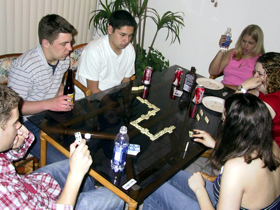

### neutral028b.bmp

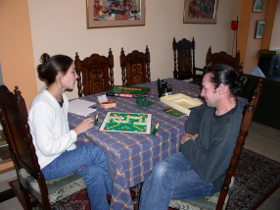

### neutral029a.bmp

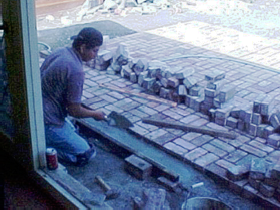

### neutral029b.bmp

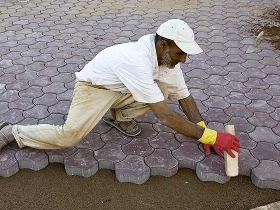

### neutral030a.bmp

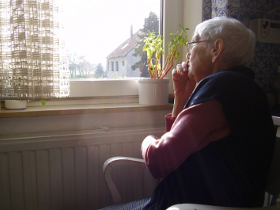

### neutral030b.bmp

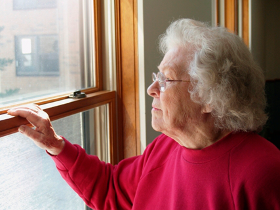

### neutral031a.bmp

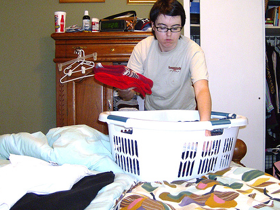

### neutral031b.bmp

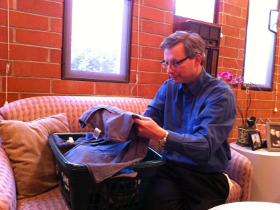

### neutral032a.bmp

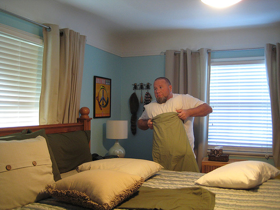

### neutral032b.bmp

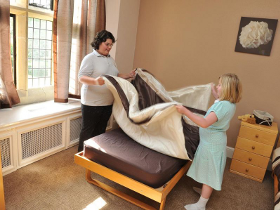

### neutral033a.bmp

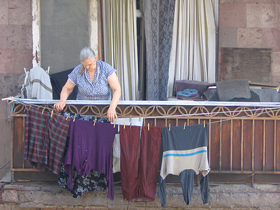

### neutral033b.bmp

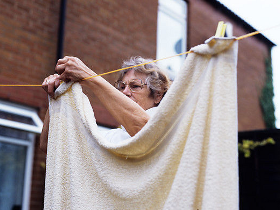

### neutral034a.bmp

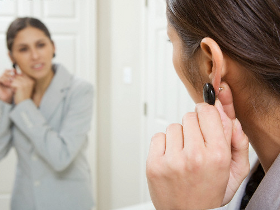

### neutral034b.bmp

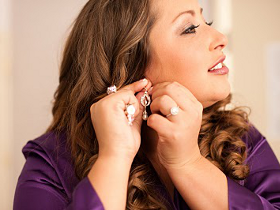

### neutral035a.bmp

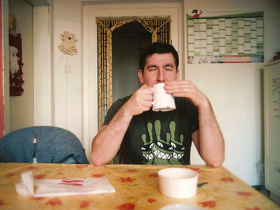

### neutral035b.bmp

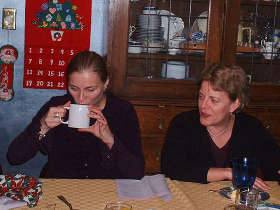

### neutral036a.bmp

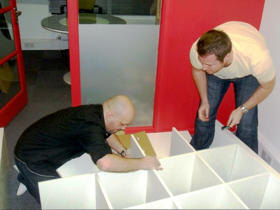

### neutral036b.bmp

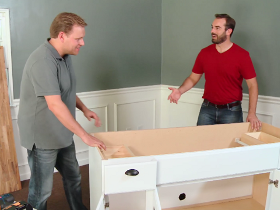

### neutral037a.bmp

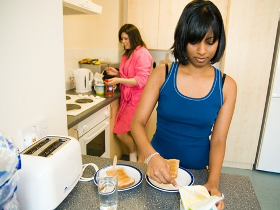

### neutral037b.bmp

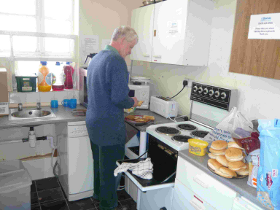

### neutral038a.bmp

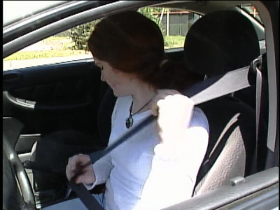

### neutral038b.bmp

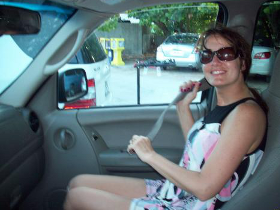

### neutral039a.bmp

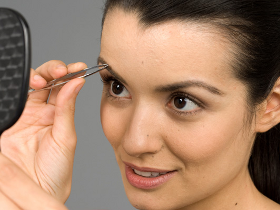

### neutral039b.bmp

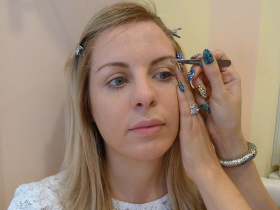

### neutral040a.bmp

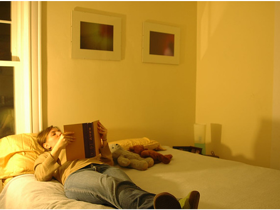

### neutral040b.bmp

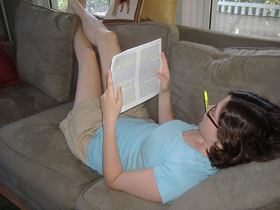

### neutral041a.bmp

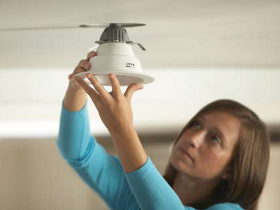

### neutral041b.bmp

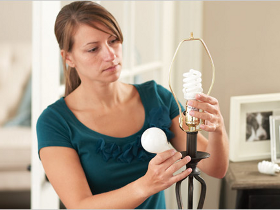

### neutral042a.bmp

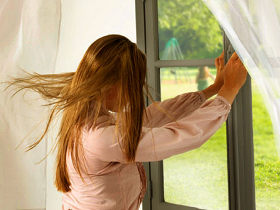

### neutral042b.bmp

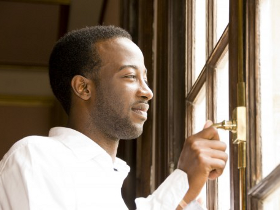

### neutral043a.bmp

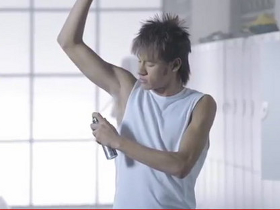

### neutral043b.bmp

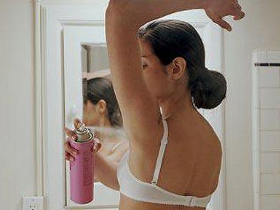

### neutral044a.bmp

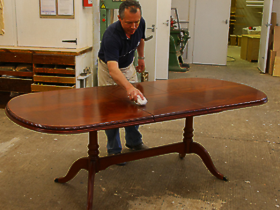

### neutral044b.bmp

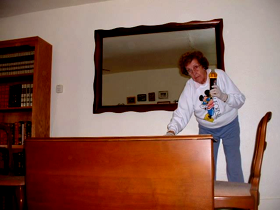

### neutral045a.bmp

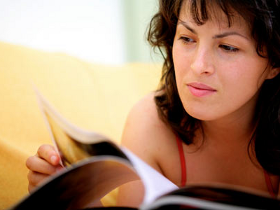

### neutral045b.bmp

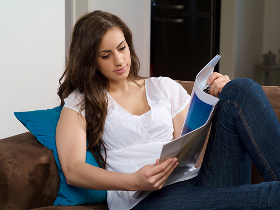

### neutral046a.bmp

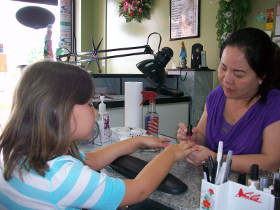

### neutral046b.bmp

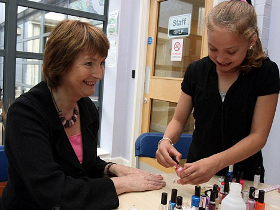

### neutral047a.bmp

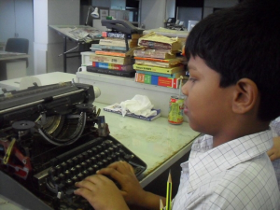

### neutral047b.bmp

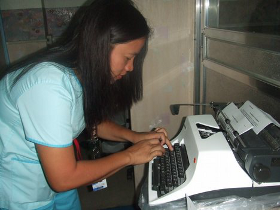

### neutral048a.bmp

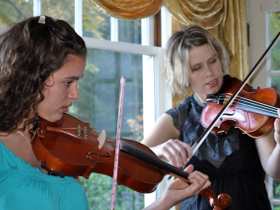

### neutral048b.bmp

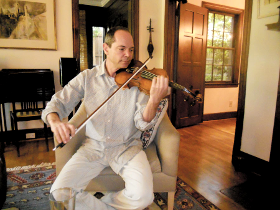

### neutral049a.bmp

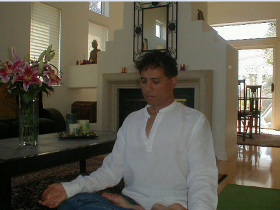

### neutral049b.bmp

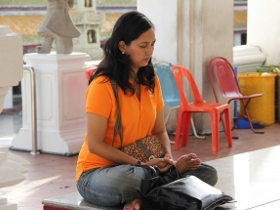

### neutral050a.bmp

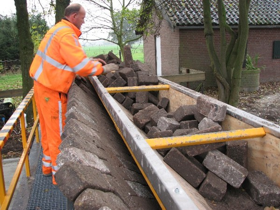

### neutral050b.bmp

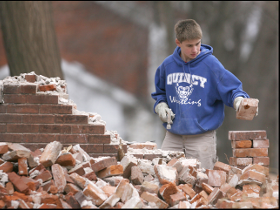

### neutral051a.bmp

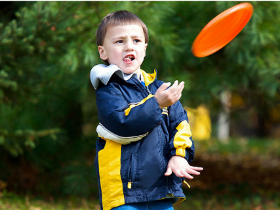

### neutral051b.bmp

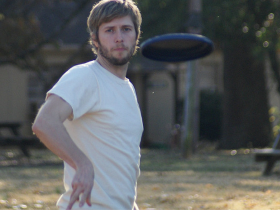

### neutral052a.bmp

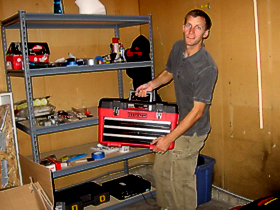

### neutral052b.bmp

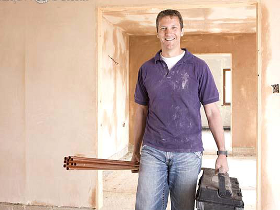

### neutral053a.bmp

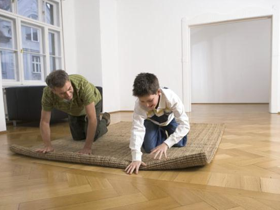

### neutral053b.bmp

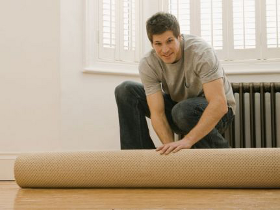

### neutral054a.bmp

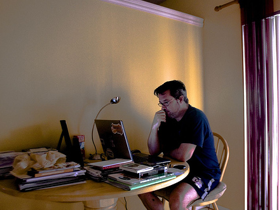

### neutral054b.bmp

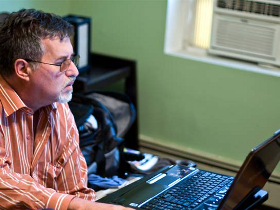

### neutral055a.bmp

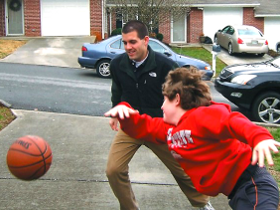

### neutral055b.bmp

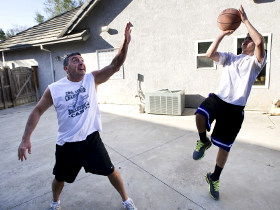
