## Supplementary figures and images for "Development of a Semantically Related Emotional and Neutral Stimulus Set"

### emotional01b.bmp

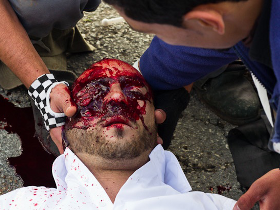

### emotional02b.bmp

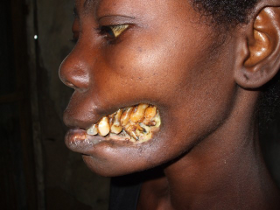

### emotional03b.bmp

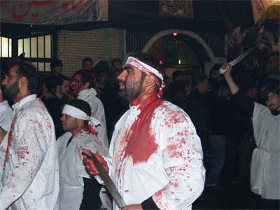

### emotional04a.bmp

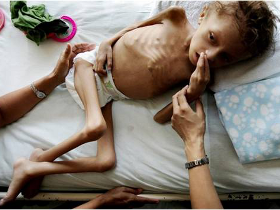

### emotional04b.bmp

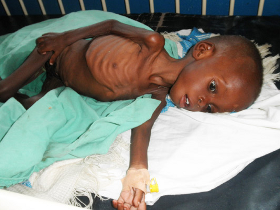

### emotional05a.bmp

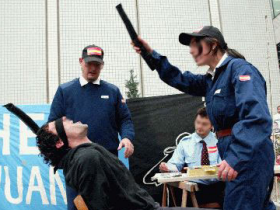

### emotional05b.bmp

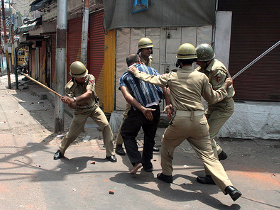

### emotional06a.bmp

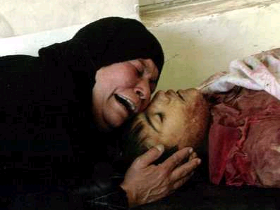

### emotional06b.bmp

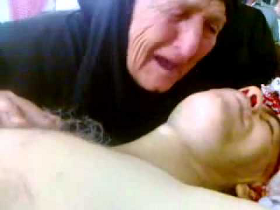

### emotional07b.bmp

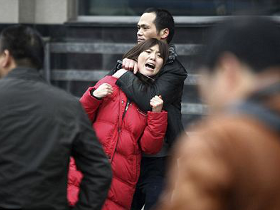

### emotional08b.bmp

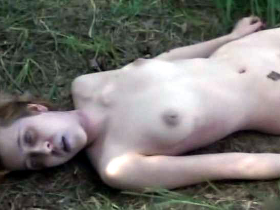

### emotional09a.bmp

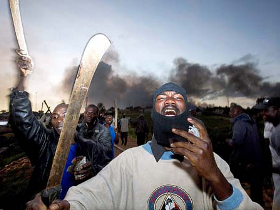

### emotional09b.bmp

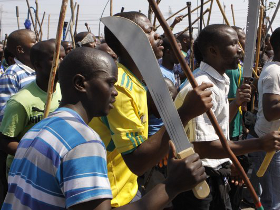

### emotional010b.bmp

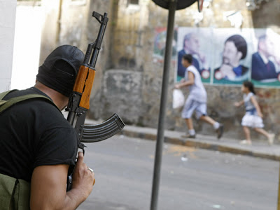

### emotional011a.bmp

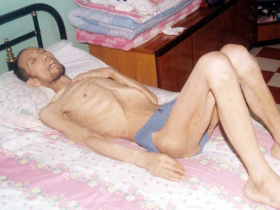

### emotional011b.bmp

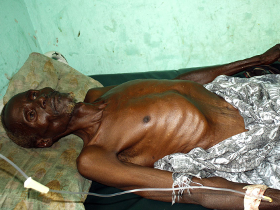

### emotional012a.bmp

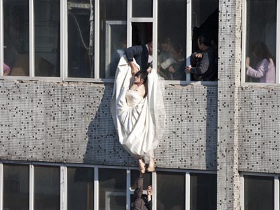

### emotional012b.bmp

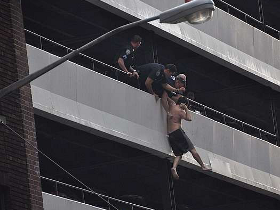

### emotional013a.bmp

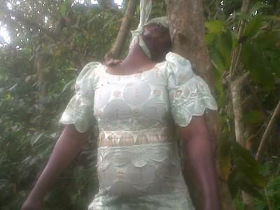

### emotional013b.bmp

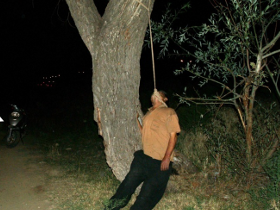

### emotional014b.bmp

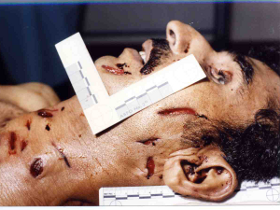

### emotional015a.bmp

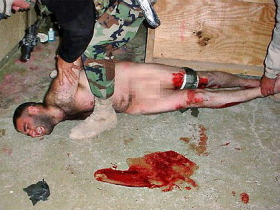

### emotional015b.bmp

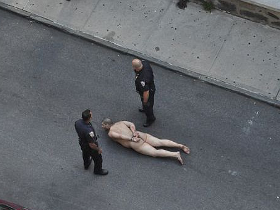

### emotional016a.bmp

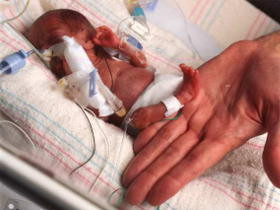

### emotional016b.bmp

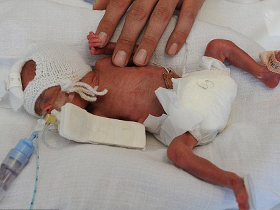

### emotional017a.bmp

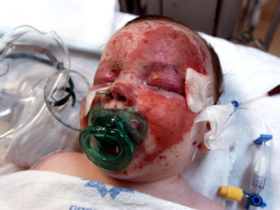

### emotional017b.bmp

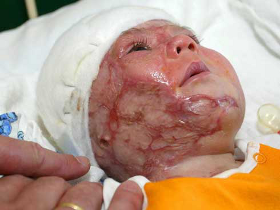

### emotional018a.bmp

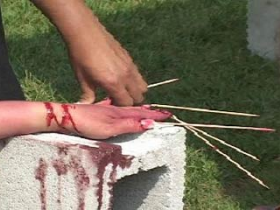

### emotional018b.bmp

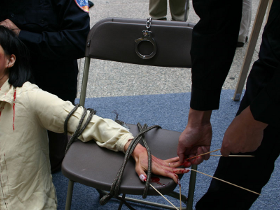

### emotional019a.bmp

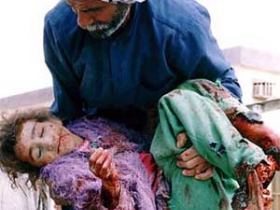

### emotional019b.bmp

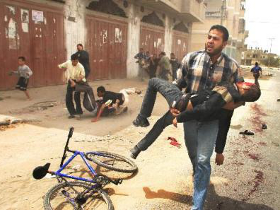

### emotional020a.bmp

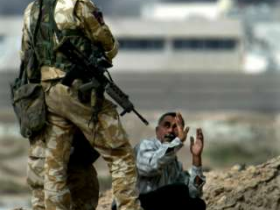

### emotional020b.bmp

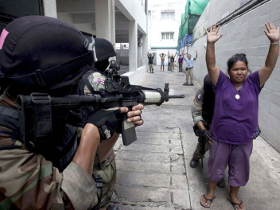

### emotional021b.bmp

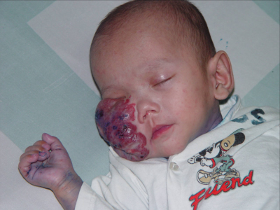

### emotional022a.bmp

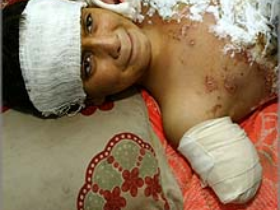

### emotional022b.bmp

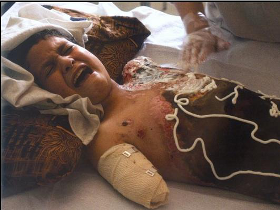

### emotional023a.bmp

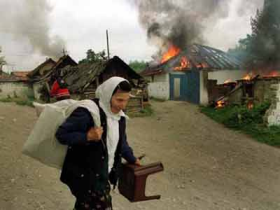

### emotional023b.bmp

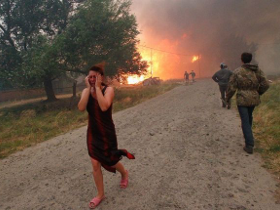

### emotional024a.bmp

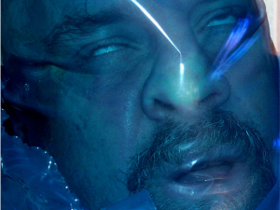

### emotional025b.bmp

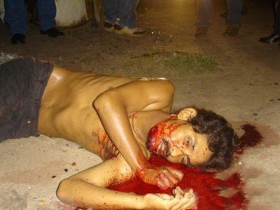

### emotional026b.bmp

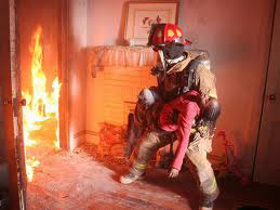

### emotional027a.bmp

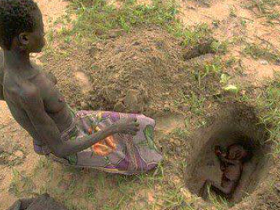

### emotional027b.bmp

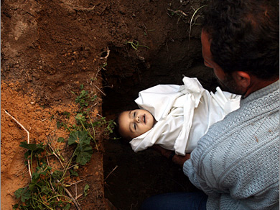

### emotional028a.bmp

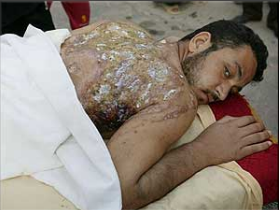

### emotional029a.bmp

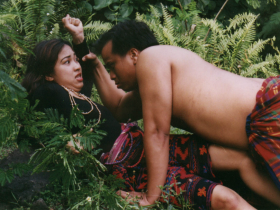

### emotional030a.bmp

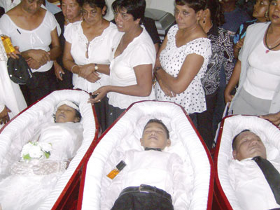

### emotional032a.bmp

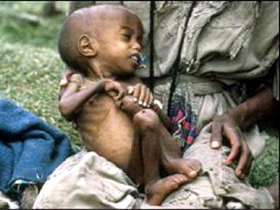

### emotional034a.bmp

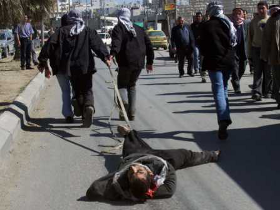

### emotional035a.bmp

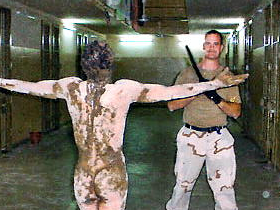

### emotional038a.bmp

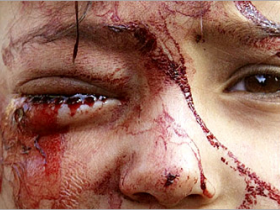

### emotional040a.bmp

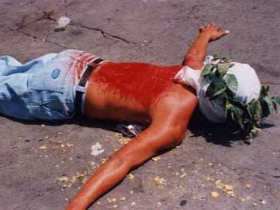

### emotional041a.bmp

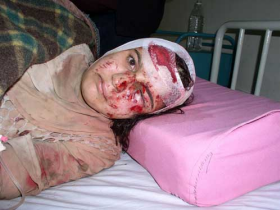

### emotional042a.bmp

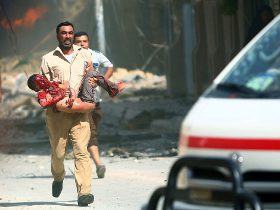

### emotional043a.bmp

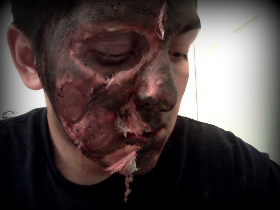

### emotional044a.bmp

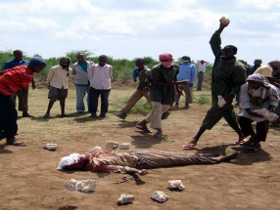

### emotional045a.bmp

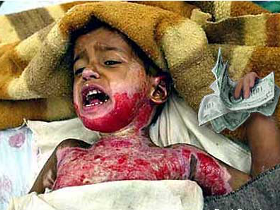

### emotional047a.bmp

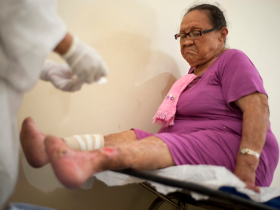

### emotional049a.bmp

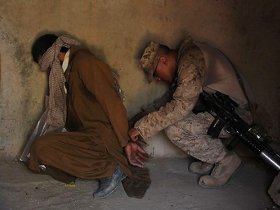

### emotional050a.bmp

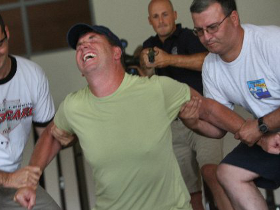

### emotional052a.bmp

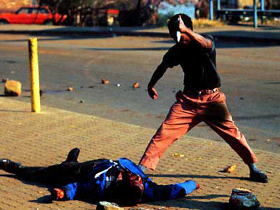

### emotional053a.bmp

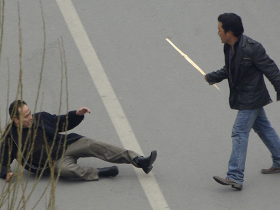

### emotional055a.bmp

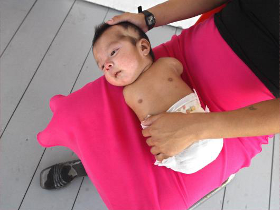

### emotional056a.bmp

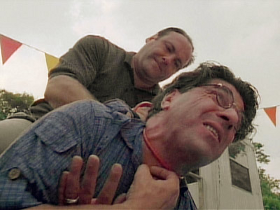

### emotional057a.bmp

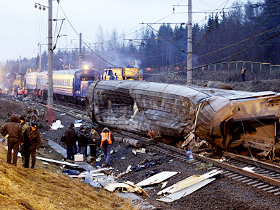

### emotional058a.bmp

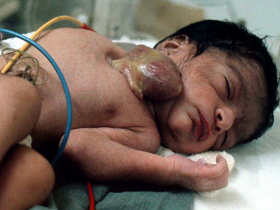

### emotional059a.bmp

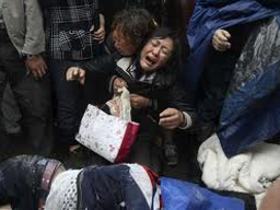

### emotional060a.bmp

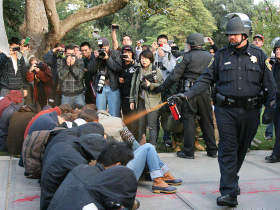

### emotional061a.bmp

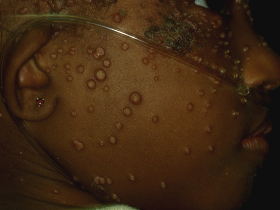

### emotional062a.bmp

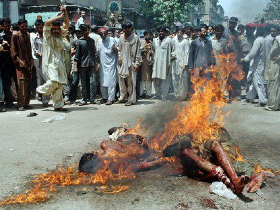

### emotional063a.bmp

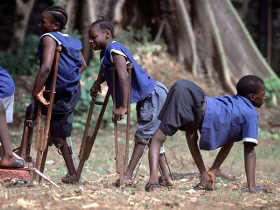

### emotional064a.bmp

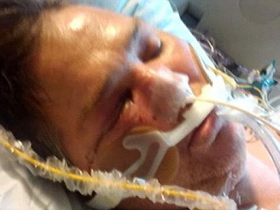

### emotional065a.bmp

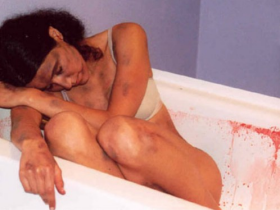

### emotional066a.bmp

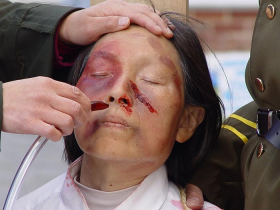

### emotional067a.bmp

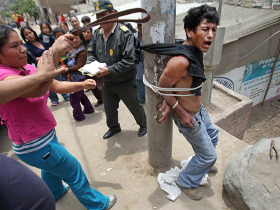

### emotional068a.bmp

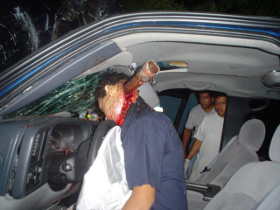

### emotional069a.bmp

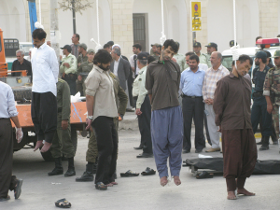

### emotional070a.bmp

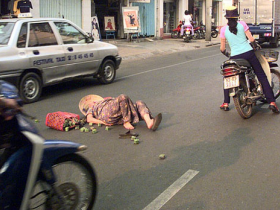

### emotional071a.bmp

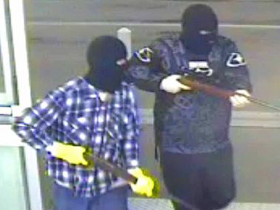

### emotional072a.bmp

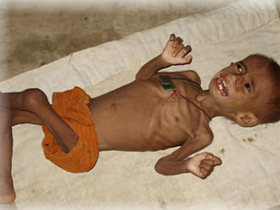

### emotional073a.bmp

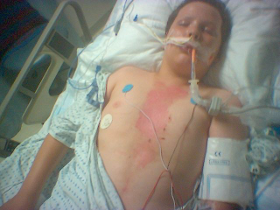

### emotional074a.bmp

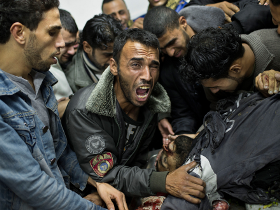

### emotional075a.bmp

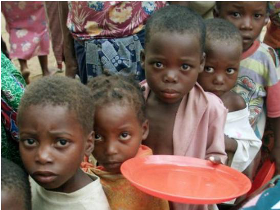

### emotional076a.bmp

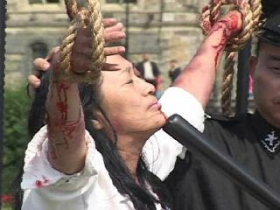

### emotional077a.bmp

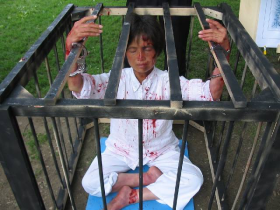

### emotional078a.bmp

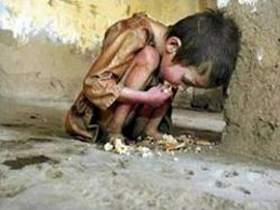

### emotional080a.bmp

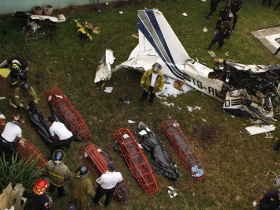

### emotional082a.bmp

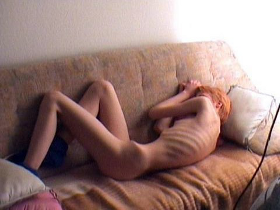

### emotional083a.bmp

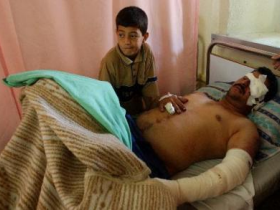

### emotional084a.bmp

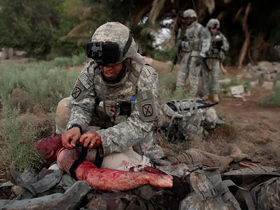

### emotional085a.bmp

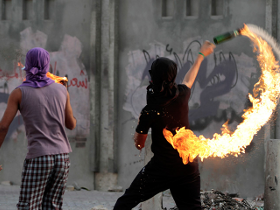

### emotional086a.bmp

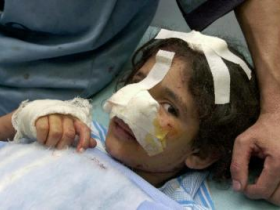

### emotional087a.bmp

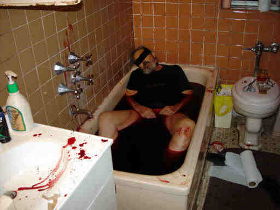

### neutral010a.bmp

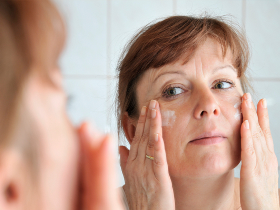

### neutral0100a.bmp

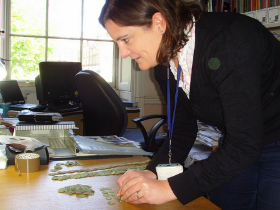

### neutral0101a.bmp

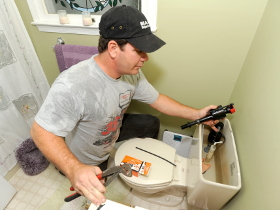

### neutral0102a.bmp

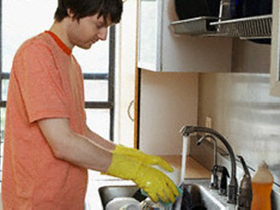

### neutral0103a.bmp

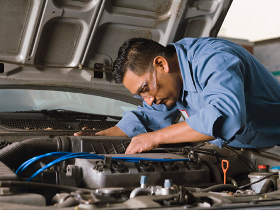

### neutral0104a.bmp

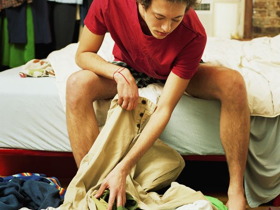

### neutral0105a.bmp

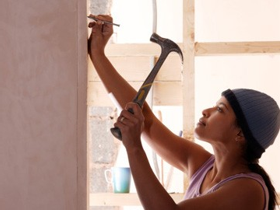

### neutral0106a.bmp

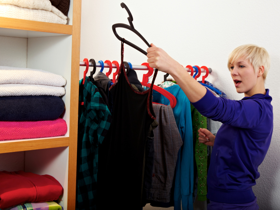
